# Defining the sediment microbiome of the Indian River Lagoon, FL, USA, an Estuary of National Significance

**DOI:** 10.1101/2020.07.07.191254

**Authors:** David J. Bradshaw, Nicholas J. Dickens, John H. Trefry, Peter J. McCarthy

**Author notes:** Corresponding author (DJB).

## Abstract

The Indian River Lagoon, located on the east coast of Florida, USA, is an Estuary of National Significance and an important economic and ecological resource. The Indian River Lagoon faces several environmental pressures, including freshwater discharges through the St. Lucie Estuary; accumulation of a anoxic, fine-grained, organic-rich sediment; and metal contamination from agriculture and marinas. Although the Indian River Lagoon has been well-studied, little is known about its microbial communities; thus, a two-year 16S amplicon sequencing study was conducted to assess the spatiotemporal changes of the sediment microbiome. In general, the Indian River Lagoon exhibited a microbiome that was consistent with other estuarine studies. Statistically different microbiomes were found between the Indian River Lagoon and St. Lucie Estuary due to changes in porewater salinity causing microbes that require salts for growth to be higher in the Indian River Lagoon. The St. Lucie Estuary exhibited more obvious microbial seasonality, such as higher Betaproteobacteriales, a freshwater associated organism, in wet season and higher Flavobacteriales in dry season samples. Distance-based linear models revealed these microbiomes were more affected by changes in total organic matter and copper than changes in temperature. Anaerobic organisms, such as Campylobacterales, were more associated with high total organic matter and copper samples while aerobic organisms, such as Nitrosopumilales, were more associated with low total organic matter and copper samples. This initial study fills the knowledge gap on the Indian River Lagoon microbiome and serves as an important baseline for possible future changes due to human impacts or environmental changes.

## Introduction

The Indian River Lagoon (IRL) is an Estuary of National Significance located on Florida’s east coast (USA) [1]. The lagoon has a total estimated annual economic value of $7.6 billion [2]. It is connected, at its southern end, to the St. Lucie Estuary (SLE), another important resource for the area [3]. The IRL has a high biodiversity because it is located at the border between temperate and sub-tropical regions, allowing it to have plant and animal species from both climates [4]. The IRL faces similar environmental issues to other estuaries, including freshwater inputs, eutrophication, organic matter, and metal contamination [1,5].

Freshwater is introduced into the IRL via runoff from local waterways and discharges from Lake Okeechobee, which are diverted into the SLE through the C-44 canal during periodic releases based upon the Lake’s water level [3]. This introduction of freshwater and its associated contaminants causes problems for the ecosystem [3] and also introduces dissolved organic material and plant matter that settles into the sediment to become the fine grained, highly organic sediment known as “IRL muck” [6]. “IRL muck” (hereinafter referred to as muck) is defined as sediment that has at least 75% water content, and the remaining solids fraction has at least 60% fines and 10% total organic matter (TOM) [7]. Muck can lead to various negative ecological impacts including nutrient flux in the water column triggering algal blooms and turbidity which damages seagrasses by blocking sunlight [8]. About 10% of the IRL is covered in muck ranging in depths from centimeters to several meters [1,7,9]. Anoxia is associated with muck and can also be caused by freshwater discharges carrying contaminants from agriculture and urban development [7,10,11]. A shift to an anoxic state alters the most energetically favorable terminal electron acceptors for microbes, altering their population, diversity, and functions [10,11]. A study of Chesapeake Bay (MD, USA), compared the microbiomes of anoxic, organic-rich, silty-clay sediments to organic-poor, sandy sediments and found major differences due to the former having microbial members that contribute to the high sulfide and methanogenic conditions in the area [12].

Other pressures on the IRL include contaminants such as trace metals which can be transported in discharges and runoff as part of metal-dissolved organic matter complexes that precipitate onto the sediment once this freshwater meets the brackish water of the IRL [6,7,13]. A survey of the northern IRL found several sites with metals above normal levels, while a survey in the SLE found a large accumulation of phosphorus and Cu, the latter likely due to Cu-containing fungicides or cuprous oxide anti-fouling paints used in marinas [14–17]. The interaction of microbes with heavy metals affects their chemical forms and therefore their solubility, mobility, bioavailability, and toxicity [18]. In turn, prokaryotic assemblages can be altered by the presence of heavy metals, which can lead to a decrease in microbial diversity and functional redundancy [5,19–22]. A recent study in Chile, found that there was a significant decrease in the abundance of bacteria in copper contaminated sites, while the abundance of archaea was similar to a less contaminated site, likely due to copper resistance mechanisms [23].

The true extent of the IRL’s biodiversity cannot be understood without information on its microbial communities [24]. Sediment microbes, especially in estuaries, face a wide range of physicochemical gradients that can cause shifts in the microbial taxonomy as well as microbial functional capabilities [13,25,26]. This study was carried out to provide the first data on the microbial communities present in the IRL and to explore potential diversity changes due to differences between: the IRL and SLE; samples with zero muck characteristics (0MC) and three muck characteristics (muck or 3MC); and non-Cu contaminated and Cu-contaminated, high TOM samples.

## Materials and methods

### Site selection

Sites were chosen based upon either being adjacent to continuous water quality monitoring stations [27,28] or known to be muck [Manatee Pocket (MP) and Harbor Branch Channel (HB)] or sandy (Jupiter Narrows and Hobe Sound) sites (Fig 1 and S1 Table). During the second year of sampling, two additional marina sites [Harbortown Marina (HT) and Vero Beach Marina] and two nearby less impacted sites (Barber Bridge and Round Island) were added. Using information from NOWData (National Weather Service), the average monthly temperature and average monthly rain sum from years 1990-2018 was obtained for the Melbourne Area, Fort Pierce Area, and Stuart 4 E Stations [29]. Streamflow data was taken from DBHYDRO (South Florida Water Management District) and from the United States Geological Services website [30,31].

**Fig 1.**
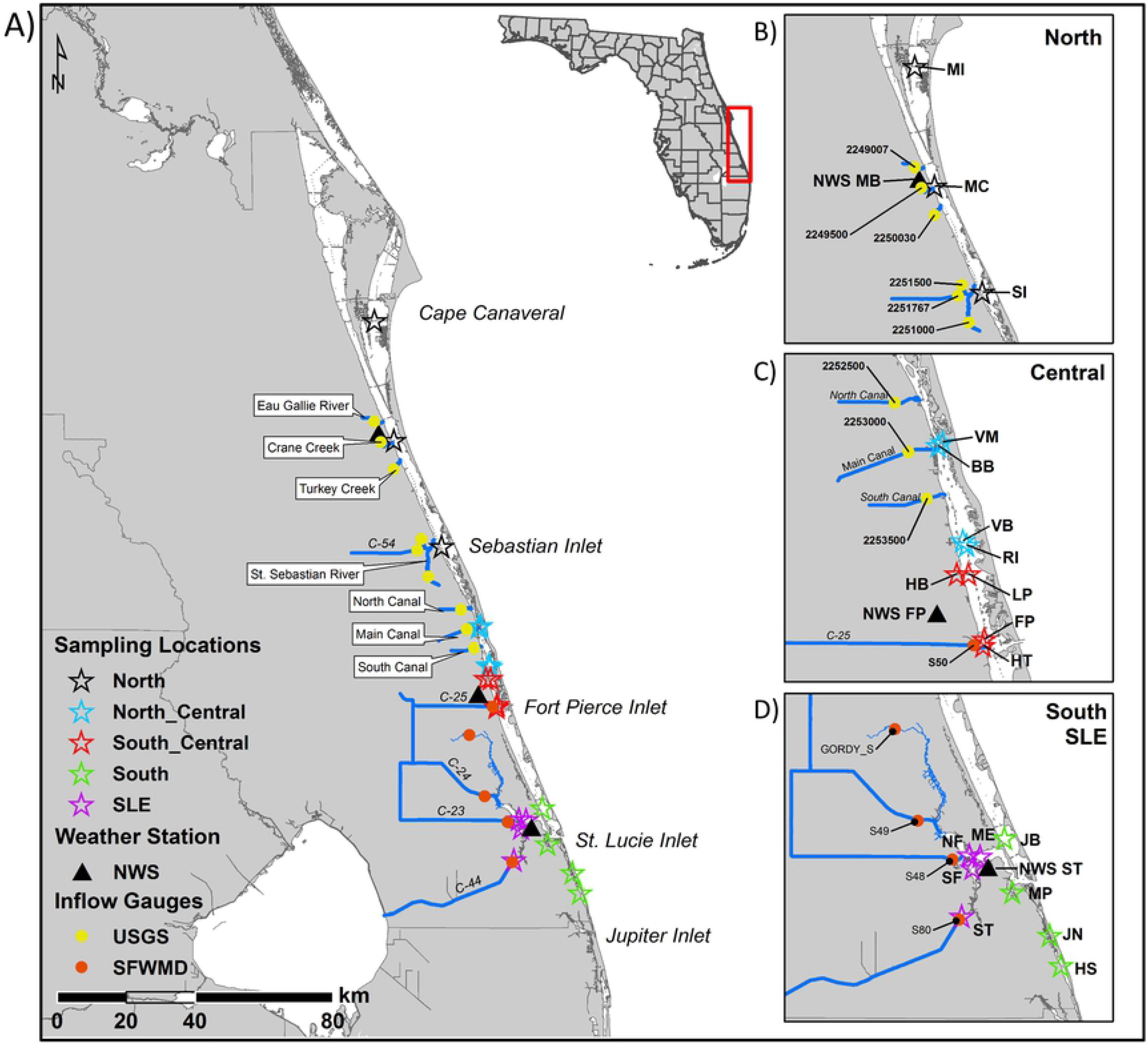
Sampling area map. The sampling area with stars indicating the site locations, triangles the location of the National Weather Service (NWS) [29] monitoring stations in Melbourne (MB), Fort Pierce (FP), and Stuart (ST), and circles the inflow gauge locations [United States Geological Service (USGS) [31] in yellow and the South Florida Water Management District (SFWMD) [30] in red]; the streams or canals associated with these gauges are denoted by blue lines. In Maps A and B black stars are the North Indian River Lagoon (IRL) sites: Merritt Island Causeway (MI), Melbourne Causeway (MC) and the Sebastian Inlet (SI). Blue stars (maps A and C) are the North Central IRL sites: Vero Beach Marina (VM), Barber Bridge (BB), Vero Beach (VB), and Round Island (RI). Red stars (maps A and C) are the South Central IRL sites: Harbor Branch Channel (HB), Linkport (LP), Fort Pierce (FP), and Harbortown Marina (HM). Green stars (maps A and D) are the South IRL sites: Jensen Beach (JB), Manatee Pocket (MP), Jupiter Narrows (JN) and Hobe Sound (HS). Maroon stars (maps A and D) are the St. Lucie Estuary (SLE) sites: North Fork (NF), South Fork (SF), Middle Estuary (ME), and South Fork 2 (ST). GPS coordinates for all study sites and environmental monitoring stations are located in S1 Table.

### Sample collection

A total of 204 sediment samples were taken during four sampling periods over a two-year period, with two sets of samples during the wet (May-Oct) and dry seasons (Nov-Apr). The samples were taken during the following months: Aug-Sep 2016 (W16), Mar-Apr 2017 (D17), Oct-Nov 2017(W17), and Apr 2018 (D18). Samples were collected with published methods [7,21]. A Wildco^®^ Ekman stainless steel bottom grab sampler was deployed from a boat to collect sediment in triplicate. The first replicate’s sediment temperature was determined with a thermometer. The top 2 cm of each replicate was subsampled with an ethanol-sterilized plastic spoon. Three sub-samples of each replicate were taken to assess the microbial community (A), dry sediment characteristics (B), and wet sediment characteristics (C). They were sealed and placed on ice until returned to the lab. Subsamples A and B were collected in sterile 50-mL Falcon tubes and 75-mL polystyrene snap-cap vials, respectively, and stored at −20° C. The remnants of the top 2 cm of the sediment were placed in a double-bagged Ziploc freezer bag as Subsample C and stored at 4° C. When ready for analysis, subsample B was thawed and dried for 48 hours at 60 °C on pre-weighed, acid-washed, glass petri dishes. The differences in pre- vs post-drying weights were used to determine water content. The dried sediment was broken up with an acid-washed mortar and pestle and sieved to remove the fraction above 2 mm (coarse) from the sample. The remaining sand (2 mm-0.063 mm) and fines (<0.063 mm) fractions were kept for further analysis. All collection plasticware was soaked in 5% HNO_3_ for a minimum of 24 hours and then rinsed three times in 18.2 MΩ deionized water.

### Metal analysis

Triplicate samples at each site were analyzed for heavy metals. Acid digests were prepared by modification of methods described in three studies [7,32,33]. Briefly, 1 g (+/− 0.0003g) of dried and sieved Subsample B was digested in 1 M HCl for one hour at 30° C with shaking at 150 rpm. Digests were filtered with DigiTubes^®^ and DigiFilters^®^ (0.45 μm) (SCP Sciences, Champlain, NY). Cu and Fe were measured with a four-point calibration curve on a Perkin Elmer 4000 atomic absorption spectrometer (Perkin Elmer, Waltham, MA). The calibration curve was rechecked every ten to twelve samples to account for absorbance drift. Reagent samples (2% HNO_3_, 1 M HCl, 18.2 MΩ deionized water) and a method control sample were analyzed to check for contamination.

### Sediment physical characteristics

A modified procedure was used to determine sediment characteristics [7]. One gram of dried and sieved Subsample B sediment was heated in a 550° C muffle furnace for four hours to burn off the organic matter. The sediment weight loss was calculated and reported as percent TOM. Grain size was determined by wet sieving 10-30 g of Sample C and drying to constant weight. The gravel, sand, and fines percentages of the total dry weight were determined. PWS was determined by centrifuging 20 g of Sample C and measuring the salinity of the resulting liquid with a portable refractometer.

### Sequence analysis

DNA was extracted from 0.25-0.3 g of sediment with the Qiagen PowerSoil DNA Isolation Kits (Hilden, Germany), and its quality checked with a Nanodrop 2000 (Oxford Technologies, Oxford, UK). Samples were sent to Research and Testing (Lubbock, TX, USA) for MiSeq 16S sequencing to amplify the bacterial/archaeal 16S V4 region with the modified primers used by the Earth Microbiome Project of 515F (GTG**Y**CAGCMGCCGCGGTAA) and 806R (GGACTAC**N**VGGGTWTCTAAT) [34,35]. The raw sequences were trimmed to remove the primers and quality-filtered with the FastX and TrimGalore programs respectively [36,37]. Quantitative Insights Into Microbial Ecology 2’s (QIIME2) [38] standard 16S workflow was used for analysis, and a Snakemake file was used for the orchestration for reproducibility [39,40]. Sequences were joined with VSEARCH [41]. Next, they were denoised with Deblur [42] run with default parameters, with the exceptions of the minimum reads parameter set to 0 to account for metadata categories with smaller sample sizes and trim length set to 232 bases. Amplicon Sequence Variants (ASVs) were annotated with a Naïve-Bayes classifier based on the scikit-learn system and the SILVA database [43,44] (version 132). Mitochondrial, chloroplast and unassigned sequences were filtered from the samples. The ASVs were aligned with MAFFT [45] and then masked [38,46] to make a phylogenetic tree with FASTTREE [47] that was then midpoint-rooted. Raw sequences have been uploaded into the National Center for Biotechnology Information Sequence Read Archive (PRJNA594146) [48].

### Statistical analysis

RStudio [49,50] (R Version 3.6.1) was used for data manipulation, visualization, generation of alpha diversity statistics (Shannon), and data manipulation. Analyses were run with the following library versions: phyloseq (1.28.0) [51], vegan (2.5-5) [52], ggplot2 (2_3.2.0) [53], reshape (0.8.8) [54], tidyverse (1.2.1) [55], and FSA (0.8.25) [56]. ASVs that did not have at least ten sequences associated with them across all samples were removed [57,58].

PRIMER7/PERMANOVA+ was also used to analyze the data [59–61]. The environmental data was checked for highly colinear variables, greater than 0.70 [62], by generating draftsman plots. This showed that TOM was positively colinear with water content, percent fines, and Fe, and negatively colinear with percent sand. This allowed TOM to represent all these variables in future analyses. The remaining environmental variables were normalized. The biological data was square root transformed, then used to make a Bray-Curtis dissimilarity matrix to create principal coordinates of analyses. Distance-based linear models were made with a stepwise selection procedure, an AICc (An Information Criterion) selection criteria, 9 999 permutations, marginal tests, and a distance-based redundancy analysis plot [21]. Overall and pair-wise permutational analysis of variance were conducted with 9 999 permutations, the unrestricted method, Type III Sum of Squares, and Monte Carlo p-values [5,63]. Overall statistical significance of environmental data and alpha diversity metrics were determined with Kruskal-Wallis or Mann Whitney U tests for categories with greater than two or just two subcategories, respectively [64,65]. Pairwise testing was conducted with the Dunn method [66]. All reported p-values were considered statistically significant if less than 0.05 after multiple testing correction with the Benjamini-Hochberg (BH) method [67]. The Snakemake file used for QIIME2 analysis and any subsequent scripts used in statistical analysis can be found on Github (https://github.com/djbradshaw2/General_16S_Amplicon_Sequencing_Analysis) [68]. Measured environmental data and metadata can be found in S2 and S3 Tables, respectively.

## Results

### Weather and streamflow discharges

Measured air temperatures were higher during each of the sampling periods than historical temperatures (1990-2018), with W16 being the hottest, followed by W17, D18, and D17 (S1 Fig). Every sampling period besides D17, which was drier than usual, was wetter than usual especially W17, for which rainfall more than doubled. Streamflow discharges matched this data with the highest streamflow occurring during W17, especially at the C44 canal leading to the South Fork 2 site (S4 Table).

### Porewater salinity and sediment temperature

Porewater salinity (PWS) and sediment temperature were measured to assess changes between sampling periods (Fig 2). Dunn testing indicated that IRL W16 and W17 sampling periods were significantly different (BH p-values < 0.05) from each other as well as both the D17 and D18 periods, although these two were significantly similar (BH p-value = 0.16) (Fig 2A and S5 Table). In the SLE, W16 and W17 were significantly similar to one another (0.65) but different from D17 and D18, which were also statistically similar to one another (0.23). The highest mean sediment temperature occurred during the W16, and Dunn testing revealed that all sampling periods were significantly different from one another except for the D17 and W17 temperatures for both the IRL (0.60) and SLE (0.052).

**Fig 2.**
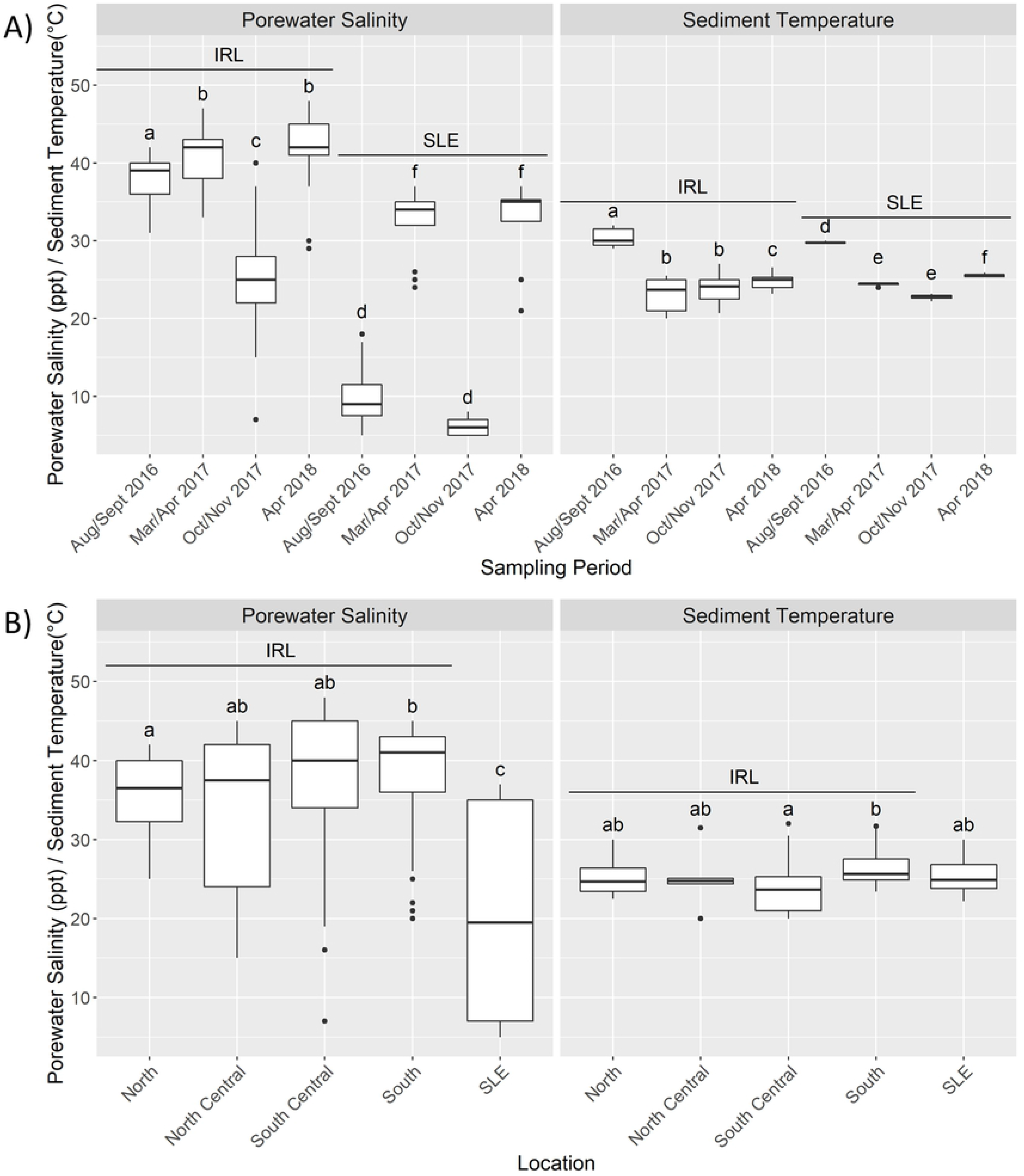
Porewater salinity and sediment temperature patterns. Porewater salinity (PWS) (left) and sediment temperature (right) by Estuary by Sampling Period (A) and by Location (B) for the Indian River Lagoon (IRL) and St. Lucie Estuary (SLE). Bars denote largest and smallest values within 1.5* the interquartile range, middle line is the median, ends of boxes are the first and third quartiles. The letters on top of each boxplot denote the results from the pairwise Dunn test with different letters denoting statistical significance (Benjamini-Hochberg adjusted p values < 0.05). In A the letters show how each of the sampling periods were different within each estuary but do not denote inter-estuary comparisons.

PWS generally increased towards the southern IRL while the SLE had the highest interquartile range, but the lowest mean (Fig 2B). All sections of the IRL were significantly different from the SLE, while only the North IRL sites were found to be statistically lower than the South IRL sites (BH p-value = 0.045) (Fig 2B and S5 Table). Sediment temperature did not vary greatly across locations, ranging from the highest mean of 26.5 °C (South) to the lowest of 24.4 °C (South Central). Each of the Location subcategories were not statistically different from one another, except for the South Central IRL being significantly lower than the South (0.00037).

### Muck and copper

The sites that, on average, exceeded three muck characteristics were Middle Estuary and South Fork, while those that exceeded at least one of the thresholds were HB, HT, Melbourne Causeway, and MP (Fig 3). None of the other 13 sites exceeded the thresholds on average. Out of the 204 samples, 40 were considered muck since their sediment characteristics exceeded three thresholds (3MC), 14 only exceeded two thresholds, 10 samples exceeded one, and 140 exceeded none (0MC) (S3 Table). 3MC samples (water content =81%, TOM = 24%, and silt/clay percentage = 81%) had 2.4x higher water content, 7.2x higher TOM, and 9.6x higher silt/clay percentages on average than 0MC samples (3.3%, 34%, 8.4%)

**Fig 3.**
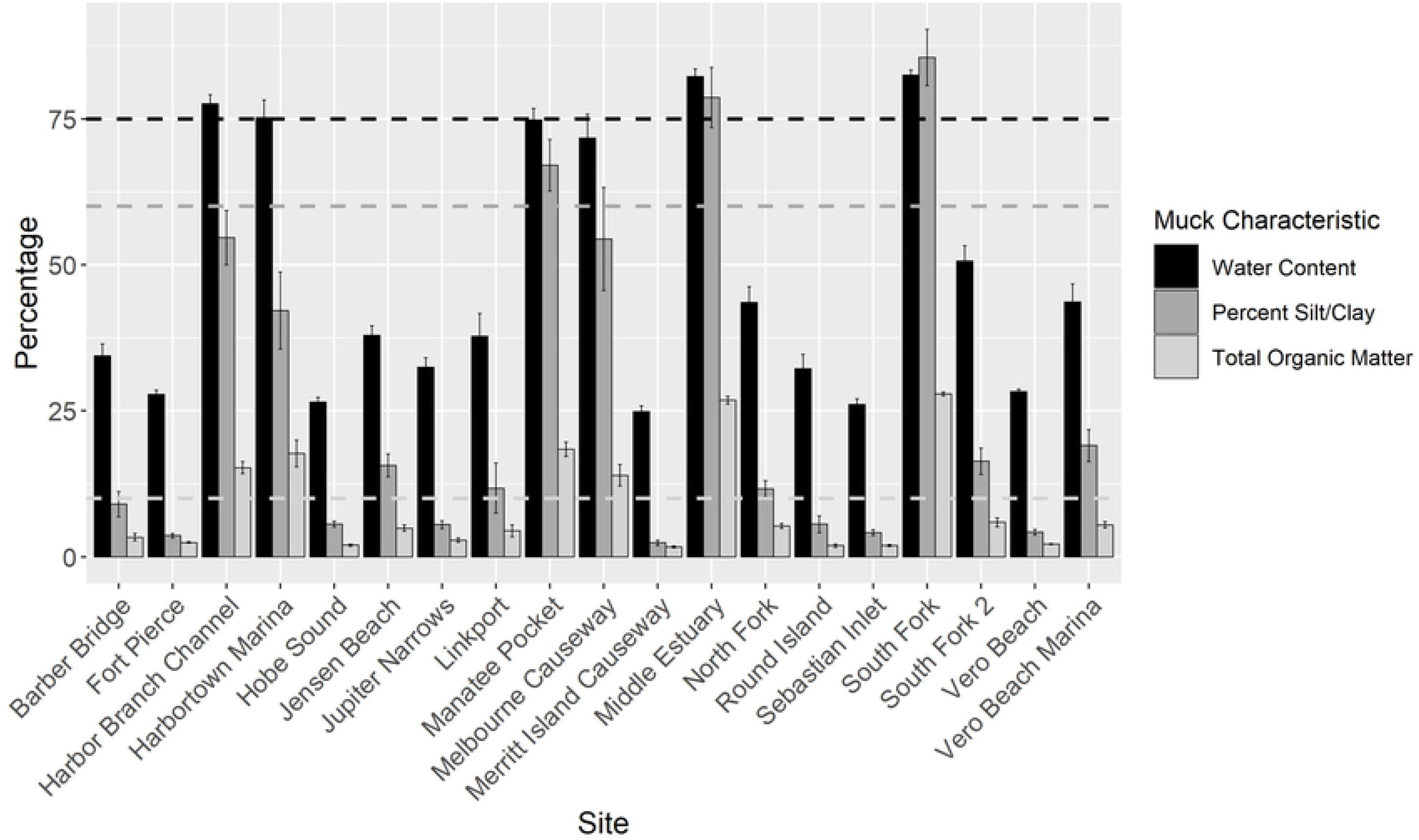
Muck characteristics by Site. Bar graph, with error bars denoting standard error, summarizing the average muck characteristics associated with each site. Black bars represent water content. The black dotted line denotes the percentage water content (75% [7]) that a site, if it exceeded all three thresholds, could be considered muck. Percent silt/clay is represented by the dark gray bars with the dark gray dotted line representing the 60% [7] threshold. Total organic matter is represented by the light gray bars with the light gray dotted line representing the 10% [7] threshold.

A sample was considered to have high TOM if it exceeded 10% and high Cu if it exceeded 65 μg/g [7,69]. Most samples that exceed at least one of the muck characteristics also had high TOM (62/64) (Fig 4 and S3 Table). The sites that had samples with both high TOM and high Cu (HiHi) included HB, MP, and HT, whereas the sites with samples that had high TOM but low Cu (HiLo) included Middle Estuary, South Fork, Melbourne Causeway, HT, Linkport, and South Fork 2. Only two samples, both from HB, were classified as having low TOM and high Cu (LoHi). The remaining 140 samples had TOM and Cu values below the thresholds (LoLo). HiHi samples (average Cu = 109 μg/g) had 3.6x and 23x more Cu than HiLo samples (30 μg/g) and LoLo samples (4.7 μg/g), respectively.

**Fig 4.**
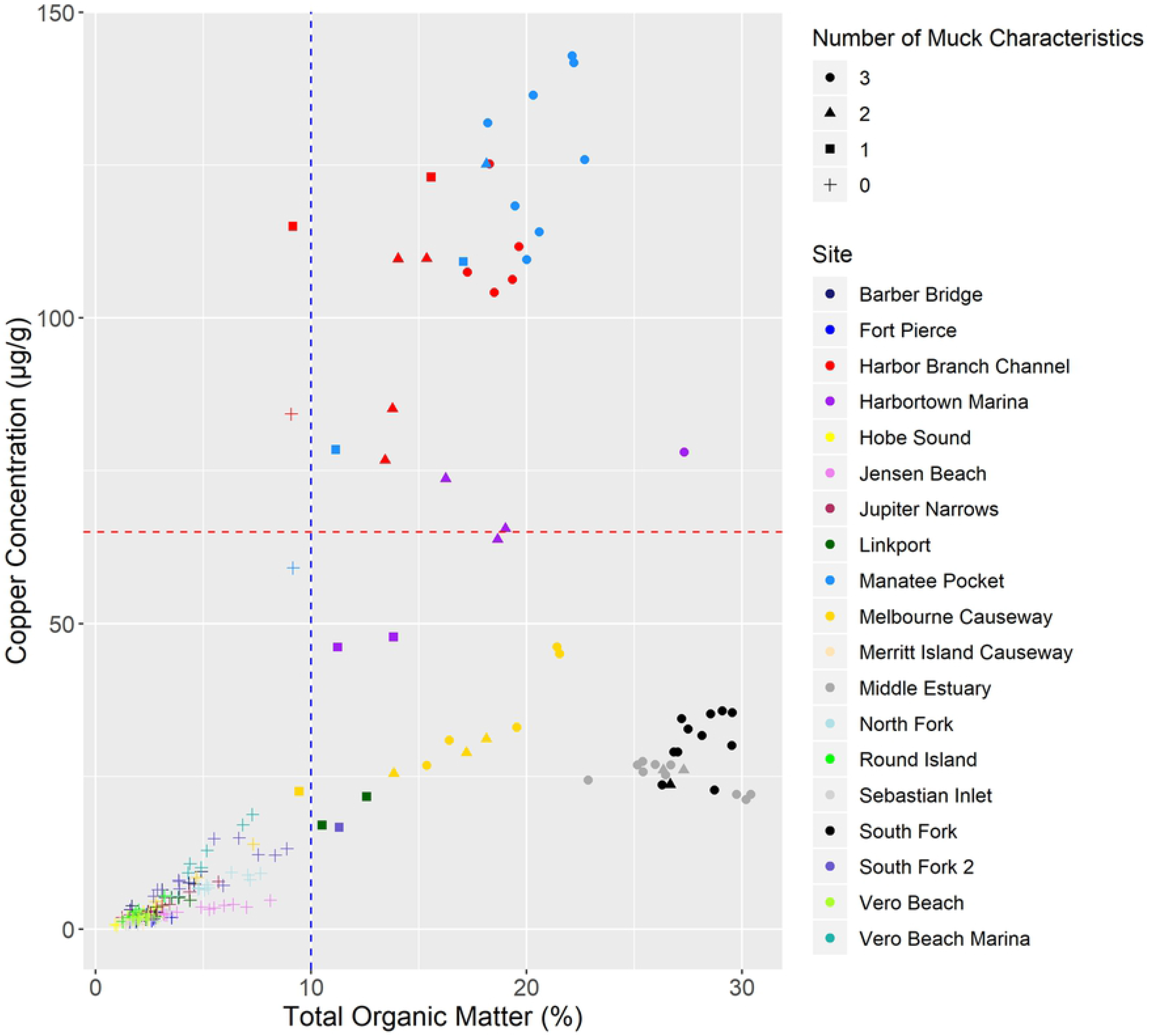
Total organic matter by copper. Point graph showing the relationship between copper (Cu) concentration (μg/g sediment) and total organic matter (TOM) percentage. Each of the 204 samples are represented by a point. Color represents the site with dark blue representing Barber Bridge, bright blue Fort Pierce, red Harbor Branch Channel, purple Harbortown Marina, yellow Hobe Sound, pink Jensen Beach, maroon Jupiter Narrows, dark green Linkport, light blue Manatee Pocket, gold Melbourne Causeway, tan Merritt Island Causeway, dark gray Middle Estuary, light blue North Fork, bright green Round Island, light gray Sebastian Inlet, black South Fork, dark purple South Fork 2, light green Vero Beach, and turquoise Vero Beach Marina. Shape represents the number of muck characteristics with circles representing three muck characteristics, triangles two, squares one, and pluses zero. The blue line at 10% [7] represents the threshold that separates the low TOM (left) from the high TOM (right) sites whereas the red line at 65 μg/g [69] separates the high Cu (above) from the low Cu (below) sites.

### General sequence information

There were 110 575 ASVs associated with the samples in this study. Filtering, described above, reduced the number of ASVs to 16 027. This filtering step also reduced the number of sequences from 1 857 744 to 1 598 653 (13.9%). The overall microbiome had 63 phyla, 193 classes, 472 orders, 799 families, 1 315 genera, and 1 691 species

### Alpha diversity

The mean Shannon alpha diversity was 6.45; its distribution was significantly correlated (p-values <2.2e-16) with observed ASVs (rho = 0.92), Fisher diversity (0.97), Simpson diversity (0.69), and Chao1 (0.92) with Spearman correlation tests. There were significant differences between Sites (BH p-value = 0.020), Estuary (0.034), Location (0.0083), Sampling Period (1.1e-15), IRL-focused Sampling Period (3.76e-15), and SLE-focused Sampling Period (3.7e-07) categories, but not by TOM/Cu (0.92), Muck (0.78), or Season (0.095) (S6 Table). Dunn analysis did not reveal any significantly different site pairs. Both the IRL and SLE exhibited the same patterns in terms of alpha diversity (Fig 5A). The D17 and D18 sampling periods were statistically similar to one another (IRL BH p-value = 0.16, SLE = 0.93); but were statistically dissimilar to the other two sampling periods, which were also significantly different from each other. Dunn testing at the Location level revealed that the North sites were statistically lower than the SLE, South, and South Central sites (Fig 5B).

**Fig 5.**
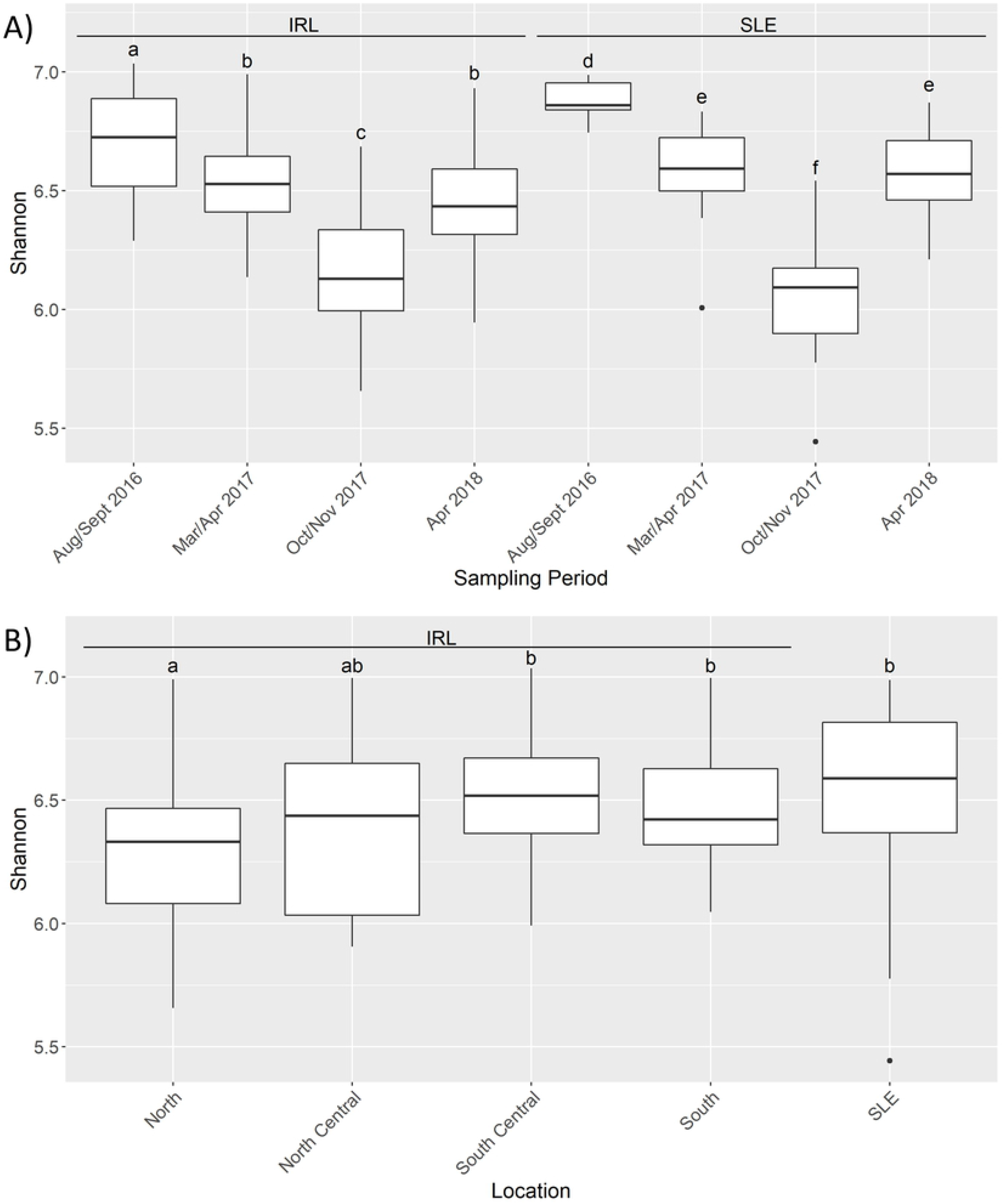
Shannon diversity paterns. Boxplots of Shannon diversity based by Estuary and Sampling Period (A) and Location (B) categories. IRL stands for Indian River Lagoon and SLE for St. Lucie Estuary. The letters on top of each boxplot denote the results from the pairwise Dunn test with different letters denoting statistical significance (Benjamini-Hochberg adjusted p values < 0.05). In A the letters show how each of the sampling periods were different within each estuary but do not denote inter-estuary comparisons. Bars denote largest and smallest values within 1.5 times the interquartile range, middle line is the median, ends of boxes are the first and third quartiles.

### Microbial community makeup of estuaries

The top three phyla in the IRL and SLE were the same: Proteobacteria, Bacteroidetes, and Chloroflexi (S2A Fig). The percentage of Epsilonbacteraeota was 16x higher in the IRL (2.2%) than the SLE (0.14%), whereas the percentage of Nitrospirae in the SLE (3.3%) was 8.7x more than in the IRL (0.38%). Desulfobacterales, Flavobacteriales, Anaerolineales and Steroidobacterales were four of the top five orders that overlapped between estuaries. (Fig 6A). The most common order for the SLE was Betaproteobacteriales (7.9%), which was 18x higher than the IRL (0.44%). The other top IRL order was Cellvibrionales (4.4%) which was 2.8x higher than in the SLE (1.5%). The following orders also occurred at levels double or greater in the IRL than in the SLE: Pirellulales (2.6x, IRL = 2.4%, SLE = 0.95%), Campylobacterales (16.0x, 2.2%, 0.14%), *B2M28* (9.6x, 1.9%, 0.20%), Actinomarinales (4.0x, 1.7%, 0.43%), and Thiotricales (3.3x, 1.7%, 0.51%). The SLE had more pronounced differences between seasons than the IRL, which had the same top five orders throughout all sampling periods. (S3 Fig) In the SLE, Betaproteobacteriales was 2.4x higher in the wet seasons (11.1%) than in the dry seasons (4.6%), whereas Flavobacteriales decreased about 4.5x between the dry (6.7%) and wet (1.5%) seasons. The following also saw decreases of at least 2x: Actinomarinales (2.1x, dry = 0.57%, wet = 0.28%), Desulfuromonadales (2.7x, 1.7%, 0.65%) Verrucomicrobiales (5.3x, 2.4%, 0.44%), Rhodobacterales (4.5x, 1.7%, 0.38%), Campylobacterales (2.8x, 0.20%, 0.072%).

**Fig 6.**
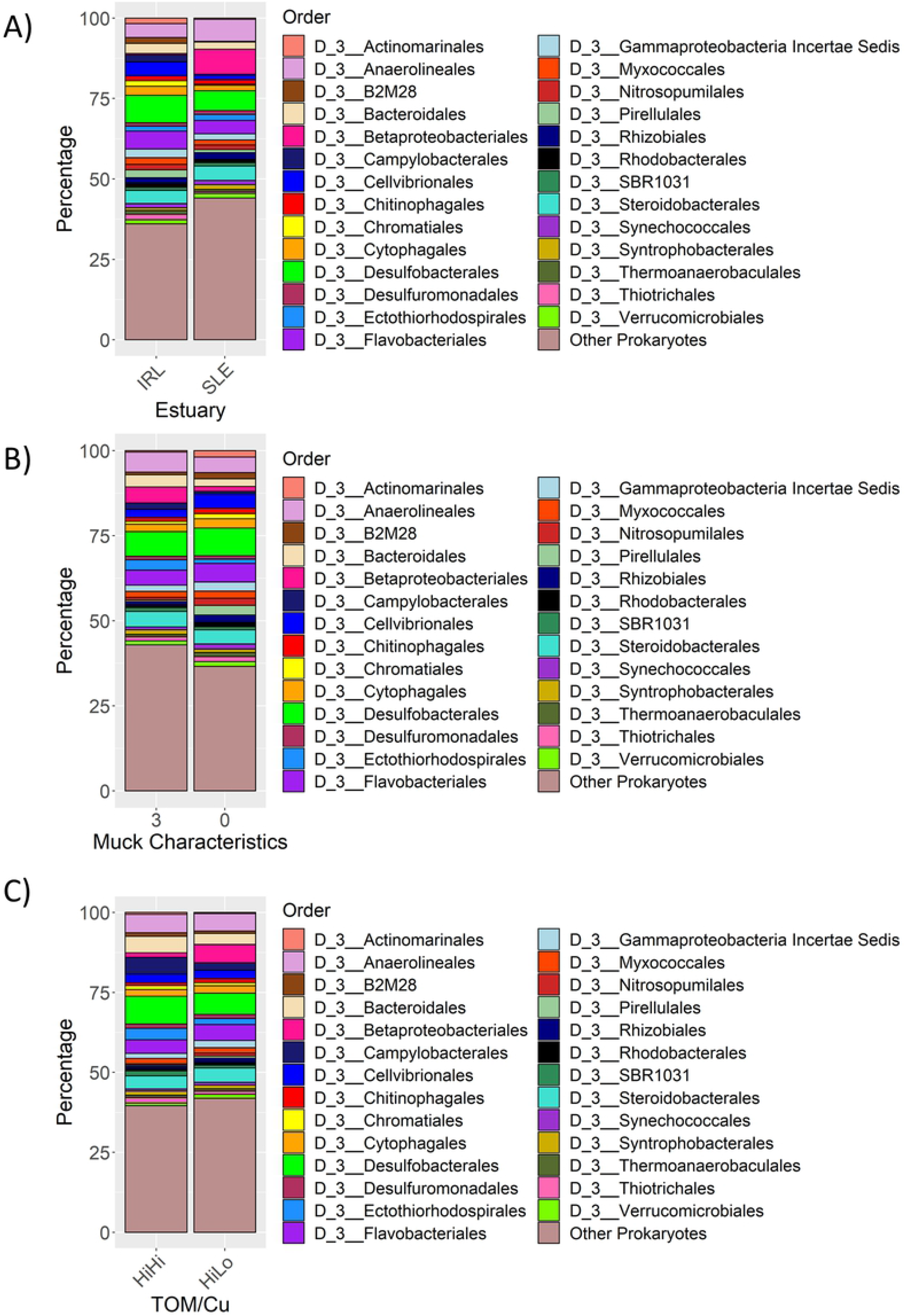
Microbial community patterns. Stacked bar graphs showing the phylogenetic orders with a mean prevalence greater than 1% across all samples associated with (A) Indian River Lagoon (IRL) and St. Lucie Estuary (SLE); (B) three muck characteristics and zero muck characteristics samples and (C) high total organic matter/high copper (HiHi) and high total organic matter/low copper (HiLo) samples. TOM stands for total organic matter and Cu stands for copper.

### Microbial community makeup of 3MC/0MC and HiHi/HiLo samples

Samples with three (3MC) and zero muck characteristics (0MC) shared four of their top five orders including Desulfobacterales, Anaerolineales, Flavobacteriales, and Steroidobacterales (Fig 6B). Bacteroidales and Cellvibrionales make up the rest of the top five for 3MC and 0MC samples respectively. Orders that were found to be at least twice as common in the 3MC samples versus the 0MC samples were: Betaproteobacteriales (3.5, 3MC = 4.8%, 0MC = 1.4%), Campylobacterales (2.3, 1.9%, 0.81%), and Ectothiorhodospirales (2.4, 3.0%, 1.3%). 0MC samples had higher levels of certain orders including Actinomarinales (4.8x, 0.39%, 1.9%), B2M28 (2.2, 0.80%, 1.8%), Nitrosopumilales (2.7, 0.76%, 2.1%), Pirellulales (5.4, 0.51%, 2.8%), Rhizobiales (2.2, 0.97%, 2.2%), and Synechococcales (2.0, 0.79%, 1.6%).

The top phyla for the high TOM and low Cu (HiLo) samples matched the order above for 3MC. High TOM and high Cu (HiHi) samples had the same top three phyla (Proteobacteria, Bacteroidetes, Chloroflexi) while Epsilonbacteraeota and Crenarchaeota replaced Acidobacteria and Planctomycetes. (S4A Fig). HiHi and HiLo shared three of the top five orders with the 3MC and 0MC samples: Desulfobacterales, Flavobacteriales, and Anaerolineales (Fig 6C). Completing the top five for HiHi was Camplyobacterales and Bacteroidales; the former was 2.2x more abundant in HiHi samples (5.2%) than in HiLo samples (2.3%). Betaproteobacteriales and Steroidobacteriales completed the top five for HiLo, with the former being found 4.0x more in HiLo samples (5.8%) than in HiHi samples (1.4%). Nitrosopumilales was 2.3x higher in the HiLo (0.91%) than the HiHi samples (0.40%).

### Beta diversity

Permutational analysis of variance results showed significant differences between the IRL and SLE samples, and across samples among the Muck and TOM/Cu subcategories (Monte Carlo p-values = 0.0001) (Table 1). 3MC and 0MC samples were statistically different from one another (0.0001), and from samples with two and one muck characteristics, which were statistically similar to one another (0.76). HiHi samples were significantly dissimilar (0.0001) from LoLo and HiLo samples but not from LoHi samples (0.32). LoLo samples were also not significantly different than LoHi samples (0.072), but were from HiLo samples (0.0001). All Location combinations were significantly different than one another (0.0001), along with most sampling periods (<0.05) except the SLE D17 and D18 sampling periods (0.29) (S7 Table).

**Table 1.** Summarized permutational analysis of variance results.

| **Overall Parameter Category<sup>a</sup>** | **Pseudo-F** | P(perms)<sup>c</sup> | P(MC)<sup>d</sup> |
| --- | --- | --- | --- |
| Pair-wise test category<sup>b</sup> | t statistic |  |  |
| **Estuary** | **42** | **0.0001** | **0.0001** |
| **TOM<sup>e</sup>/Cu<sup>f</sup>** | **7.2** | **0.0001** | **0.0001** |
| Low TOM/Low Cu, High TOM/High Cu | 2.7 | 0.0001 | 0.0001 |
| Low TOM/Low Cu, Low TOM/High Cu | 1.2 | 0.045 | 0.072 |
| Low TOM/Low Cu, High TOM/Low Cu | 3.7 | 0.0001 | 0.0001 |
| High TOM/High Cu, Low TOM/High Cu | 1.1 | 0.47 | 0.32 |
| High TOM/High Cu, High TOM/Low Cu | 2.8 | 0.0001 | 0.0001 |
| Low TOM/High Cu, High TOM/Low Cu | 1.4 | 0.034 | 0.061 |
| **Muck Characteristics** | **5.4** | **0.0001** | **0.0001** |
| 0, 2 | 1.9 | 0.0002 | 0.0001 |
| 0, 3 | 3.4 | 0.0001 | 0.0001 |
| 0, 1 | 1.6 | 0.0028 | 0.0022 |
| 2, 3 | 1.4 | 0.03 | 0.034 |
| 2, 1 | 0.82 | 0.88 | 0.76 |
| 3, 1 | 1.5 | 0.013 | 0.017 |
| **Estuary by Season** | **17** | **0.0001** | **0.0001** |
| All pairwise analyses had P(MC) values equal to 0.0001<sup>g</sup> |  |  |  |
| **Location** | **10** | **0.0001** | **0.0001** |
| All pairwise analyses had P(MC) values equal to 0.0001<sup>g</sup> |  |  |  |
| **Estuary by Sampling Period** | **10** | **0.0001** | **0.0001** |
| SLE-D17, SLE-D18 | 1.1 | 0.25 | 0.29 |
| All other pairwise analyses had P(MC) values less than 0.05<sup>g</sup> |  |  |  |
| **Site** | **11** | **0.0001** | **0.0001** |
| Barber Bridge, Vero Beach Marina | 1 | 0.31 | 0.4 |
| All other site pairwise analysis have P(MC) values below 0.05<sup>g</sup> |  |  |  |

<sup>a</sup>Bold text represents results from overall category and <sup>b</sup>regular text represents results from the pair-wise testing results.
<sup>c</sup>P(perms) stands for the permutation p value, <sup>d</sup>P(MC) stands for Monte Carlo p value, <sup>e</sup>TOM stands for total organic matter, and <sup>f</sup>Cu stands for copper. <sup>g</sup>The full results can be seen in S7 Table.

### Influence of environmental variables

The environmental variable most statistically associated with microbial variation between the samples was PWS (Pseudo-F = 20.396, proportion =0.09171), followed by TOM (15.244, 0.064028), Cu (7.5017, 0.030522), and finally sediment temperature (5.5491, 0.022076) (Fig 7 and S8 Table). PWS generally decreased from the upper left corner to lower right, separating the IRL sites from the SLE sites. TOM generally decreased from the lower left corner to the upper right, separating the samples with muck characteristics from those with none. Cu generally decreased from the top of the graph, where the HiHi and LoHi samples were found, to the bottom, where the HiLo and LoLo samples were.

**Fig 7.**
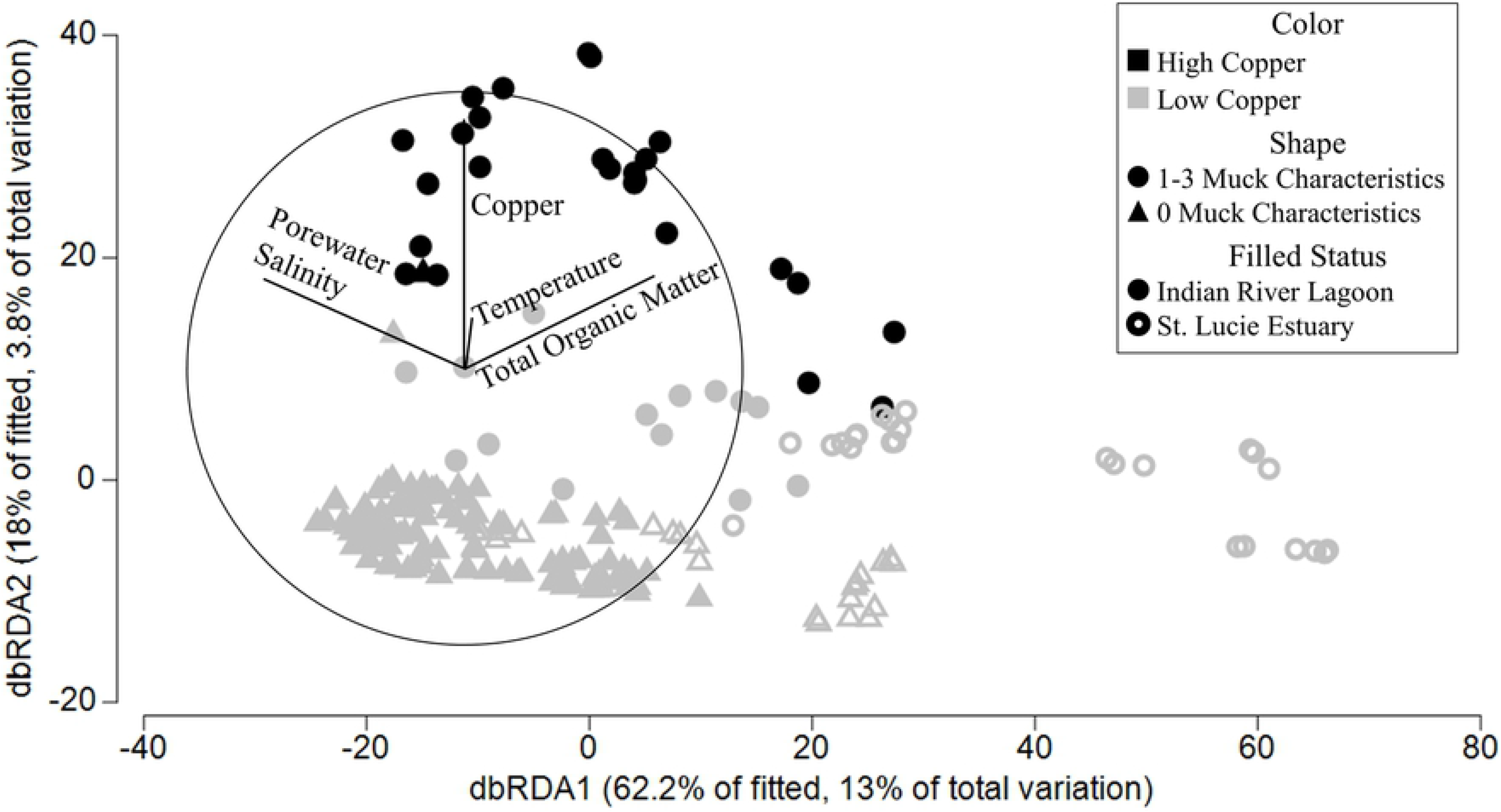
Distance-based redundancy analysis. Distance-based redundancy analysis of the sediment samples with colors representing the amount of copper [high or > 65μg/g [69] (black), low or < 65μg/g (gray)]. The shape shows the number of muck characteristics associated with a sample with circles representing 1-3 characteristics and triangles representing no muck characteristics. A filled shape is an Indian River Lagoon sample and a hollow shape represents a St. Lucie Estuary sample. The results of the distance-based linear models are shown by the lines and their associated environmental parameter and shown in S8 Table.

## Discussion

### Environmental parameters and seasonality

Sampling was scheduled to capture seasonality, but the extended impacts of Hurricane Irma which made landfall in Florida, USA on September 10, 2017 shifted one of the sampling periods from Aug-Sept 2017 to Oct-Nov 2017. PWS seasonality was more defined in the SLE than the IRL since there was a statistically lower PWS in the SLE during both wet seasons compared with the dry seasons (Fig 2A). This reflects the increased rainfall and consequent discharges from Lake Okeechobee/C44 and other canals/streams during these times (S1 Fig and S4 Table). In contrast, the PWS in the IRL was statistically different between the wet seasons (Fig 2A). Although the sediment temperature was highest during W16, W17 was either statistically similar to or lower than the two dry seasons in both the IRL and SLE (Fig 2A). In conclusion, seasonality was a prominent environmental factor in the SLE in terms of PWS but not sediment temperature; however, neither parameter showed distinguishable seasonality in the IRL. PWS generally increased from North to South in the IRL, possibly due a greater number of oceanic inlets in the south (Figs 1 and 2B) [4,70]. Owing to higher freshwater releases, the SLE had a statistically lower PWS with a wide range (Fig 2B and S4 Table) [70].

### Muck accumulation

Muck is formed by the bacterial degradation of organic matter at the transition between freshwater and estuarine waters [6,7]. The sites that had at least one muck sample were located near this transition in either the SLE (South Fork, Middle Estuary) or C25 (HT) (Figs 1 and 3). Samples with at least one muck characteristic were also near areas that allowed accumulation of organic matter due to restricted flow caused by the shape and bathymetry of the area (HB, MP) or high residence time (MC) (Figs 1 and 3) [71,72]. Although South Fork 2 is located adjacent to the C44 canal, it experiences periodic large volumes of high velocity flow that could prevent fine particles and organic matter from settling (Fig 1).

### Copper contamination

Copper can be found in the sediment near marinas due to the use of copper-antifouling paints, which explains why HB, located in a channel historically used for large boats, HT, located by a large active marina, and MP, located by a small active marina, were the only sites to have high Cu (Figs 1 and 4) [69,73]. The site by the active Vero Beach Marina is not located in a flow-restricted area, possibly allowing the current to move contaminants away from the area (Fig 1).

### Alpha diversity patterns

Alpha diversity was highest in the warmest period (W16) and increased from north to south, matching the pattern of higher diversity in warmer environments seen in other studies (Figs 5A and 5B) [74]. The diversity drop associated with W17 samples may be related to the impact of Hurricane Irma (Fig 5A) [75].

### Estuarine microbial community differences

PWS has been identified as an important factor in microbiome variation in other studies [63]. The most common phylum associated with either estuary was Proteobacteria, which is consistent with other estuarine studies (S2A Fig) [12,63,75,76]. Proteobacteria has members that are capable of utilizing a wide variety of substrates, which allow them to occupy many different environments [77]. The most common Proteobacteria orders in the IRL and SLE included Desulfobacterales and Steroidobacterales, while the IRL had more Cellvibrionales, and the SLE had more Betaproteobacteriales (Fig 6A). Most members of Desulfobacterales, including its high-percentage families Desulfobacteraceae and Desulfobulbaceae (S2C Fig), are sulfate-reducing bacteria and have also been found in high percentages in other estuary studies [63,78]. The family Woeseiaceae (order Steroidobacterales) (S2C Fig) has members capable of facultative sulfur- and hydrogen-based chemolithoautotrophy and is considered a core member of marine sediments [79,80]. The family Halieaceae (order Cellvibrionales) (S2C Fig) is found in coastal marine areas and is capable of aerobic photoheterotrophic growth [81]. *Halioglobus* (S2D Fig) (Halieaceae) is capable of denitrification and requires NaCl for growth, which may be why it was more prevalent in the IRL [82]. Betaproteobacteriales was shown in multiple studies to increase in freshwater-influenced areas of estuarine lagoons, which is similar to this study where Betaproteobacteriales increased in the SLE during the wet season samples (Fig 6A and S3 Fig) [63,76]. Flavobacteriales (phylum Bacteroidetes) (Fig 6A) is also commonly abundant in other estuary studies [75]. Most members of the family Flavobacteriaceae (S2C Fig) require NaCl or seawater salts for growth, which explains the decrease in Flavobacteriales during the wet seasons in the SLE (S3 Fig) [83]. Anaerolineaceae (phylum Chloroflexi, order Anaerolineales) (Fig 6A and S2A Fig) is comprised of obligate anaerobes with most members capable of breaking down proteinaceous carbon sources [84]. Behera et al. (2017), in another study into the effects of freshwater on a brackish lagoon, found that the phylum Acidobacteria and classes Gammaproteobacteria and Alphaproteobacteria were higher in more marine environments [63]. Our study instead found that there were more Acidobacteria in the more freshwater SLE along with relatively equal amounts of Gamma-(0.61% more in IRL) and Alphaproteobacteria (0.13% more in SLE) (S2A and S2B Figs).

Microbes that are highly abundant in a system are likely to be the actively-metabolizing part of the community; although a portion of the rare microbes can be active, they are more likely to be dormant or dead cells [85,86]. Since Desulfobacterales and Steroidobacterales had high relative abundances, they were likely responsible for some of the sulfur cycling in the lagoon [78,80]. Likewise, carbon cycling was likely affected by the photosynthetic Cellvibrionales and members of Anaerolineales [82,84]. Nitrogen cycling also was affected by denitrifying members of Cellvibrionales, like Halioglobus and nitrogen-fixing members of Betaproteobacteriales [78,81].

### Microbiome shifts associated with copper and muck

Alpha diversity did not decrease in stressed sediments (sediment with muck characteristics or Cu-contamination) as seen in other studies exploring the effects of metals and clay/silt [5,19]. While a diversity decrease can be an indicator of impaired environmental health, organisms can also become adapted to stressors with the largest drop in diversity associated with initial exposure to contamination [5]. Thus, it is possible the community has had enough time to adapt to contamination and for tolerant species to flourish [87]. There were significant differences between the 0MC and 3MC microbiomes, which could be partially due to the smaller pore size in muck affecting the ability of some microbes to flourish [88].

TOM has also been seen as an important environmental variable in other studies [89] or studies that measured TOM-covariable parameters such as percent fines [5] or silt [21]. The top phyla matched between the 0MC and IRL samples as well as the 3MC and SLE samples (S2A and S4A Figs). This pattern likely occurred because 43.8% (21/48) of the samples in the SLE were classified as 3MC samples whereas only 12.2% (19/156) of the IRL sites were classified as 3MC samples. This could help explain why there were higher percentages of Betaproteobacteriales in the 3MC samples as well as the HiLo samples since there were also no HiHi samples in the SLE samples.

In comparison to HiLo samples and 0MC samples, HiHi and 3MC samples had higher percentages of Epsilonbacteraeota and Crenarchaeota (S4A and S5A Figs). A recent study suggested that Archaea, such as the Crenarcheota, have a greater resistance to copper contamination due to their ability to sequestrate or pump out copper; this study also found Crenarchaeota flourishing in their copper-contaminated samples [23]. Members of the order Campylobacterales (phylum Epsilonbacteraeota) were also found at higher abundances in HiHi and 3MC sediments; some of its members, including the genus *Sulfurovum*, have been found in sulfide- and hydrocarbon-rich sediments similar to muck [90] as well as metal-contaminated sediments [91,92]. (Figs 6B and 6A; S4D and S5D Figs). *Sulfurovum* is a mesophilic facultative anaerobe, requires salts for chemolithoautotrophic growth with elemental sulfur or thiosulfate as an electron donor, nitrate or oxygen as an electron acceptor, and CO_2_ as its carbon source [93]. Conversely, 0MC and HiLo samples had higher abundances of the *Candidatus Nitrosopumilus* genus and its associated higher taxonomic ranks *Nitrosopumilus* is similar to *Sulfurovum* in that it uses CO_2_ as its carbon source and is halophilic, but it grows chemolithoautotrophically by conducting ammonia oxidation to nitrite and is aerobic. Other families which are typically aerobic and were more abundant in the 0MC samples included Pirellulaceae and Sandaracinaceae [94–96]. This shows that the microbiome differences between 3MC and 0MC samples were at least partially due to the former being typically more anaerobic since increased organic matter can lead to increased respiration and depletion of oxygen [7]. 3MC samples also had lower abundances of Cyanobiaceae which could be due to the increased turbidity associated with muck and its higher percentage of silt/clay [7]. Sediment microbial communities have been shown in other studies to be greatly affected by carbon sources, electron acceptors, and amount of oxygen in an area [12,89].

## Conclusions

The most important variable causing shifts between the microbiomes was PWS, this was mainly due to the influence of seasonal freshwater discharges into SLE causing microbiome differences in comparison to the IRL. Other observed differences included increases in anaerobic organisms in the higher TOM 3MC samples and aerobic organisms in the lower TOM 0MC samples. Tracking changes in the differentially abundant microbes present in different sediment types will allow management agencies to predict areas that are at risk of developing muck due to microbial influences or becoming sufficiently copper-contaminated to cause biological harm. This study provides the first NGS data on the microbial diversity of the IRL which will serve as an important baseline for future studies to measure the impact of anthropogenic inputs and natural disasters. This data can also be used by researchers in other estuarine areas to compare their results to determine if their systems are facing similar shifts in the microbiomes due to similar anthropogenic impacts.

Future studies should be performed with greater sequencing depth and higher sampling frequency, which could allow more of the diversity and rarer taxa in the samples to be captured, and shotgun metagenomics to identify functional differences between sites. Incorporating the measurement of anoxia and biogeochemical cycles would help to further delineate which environmental variables are causing shifts to the microbiomes between sediment types and geographical locations.

## Acknowledgements

We thank Dennis Hanisak and Joshua Voss for their guidance throughout this project; Gabrielle Barbarite, Austin Fox, Stacey Fox, Dedra Harmody, John Hart, Hunter Hines, Brandon McHenry, and Carlie Perricone for their advice and contributions to the research. A special thanks to Emily Sniegowski for her support and thoughtful edits.

## Supporting information

**S1 Fig. National Weather Service temperature and rain sum patterns.** National Weather Service NOWData showing the four sampling periods (Aug-Sep 2016 (dark orange), Mar-Apr 2017a (dark green), Oct-Nov 2017b (light orange), Apr 2018 (light green)) as well as the historical temperature (A) and rain sum (B) during those months for the years 1990-2018 (dark gray). Bars denote standard error.

**S2 Fig. Other taxonomic levels by Estuary.** Stacked bar graphs showing the phyla (A), classes (B), orders (C), families (D), and genera (E) that have a mean of greater than 1% across all samples. These graphs show the differences between the two main basins of the study, (Indian River Lagoon (IRL) or St. Lucie Estuary (SLE))

**S3 Fig. Estuary orders by Sampling Period.** Stacked bar graph showing the orders with a mean greater than 1% across all samples grouped by estuary (Indian River Lagoon (IRL) or St. Lucie Estuary (SLE)) and sampling period (Aug/Sept 2016, Mar/Apr 2017, Oct/Nov 2017, and Apr 2018).

**S4 Fig. Other taxonomic levels by Muck classification.** Stacked bar graphs showing the phyla (A), classes (B), orders (C), families (D), and genera (E) that have a mean of greater than 1% across all samples. These graphs show the differences between the samples with three and zero muck characteristics.

**S5 Fig. Other taxonomic levels by Total Organic Matter/Copper classification.** Other taxonomic levels by Total Organic Matter/Copper classification Stacked bar graphs showing the phyla (A), classes (B), orders (C), families (D), and genera (E) that have a mean of greater than 1% across all samples. These graphs show the differences between the samples with high TOM and copper (HiHi) and high TOM and low copper (HiLo).

**S1 Table. GPS Coordinates.** ^a^NWS stands for National Weather Service [29], ^b^IRFWCD for Indian River Farms Water Control District, ^c^USGS for United States Geological Service [31], ^d^SFWMD for South Florida Water Management District [30].

**S2 Table. Measured environmental variables per sample.** Sediment temperature was determined using a thermometer, water content by weight loss during oven drying, total organic matter by weight loss in a muffle furnace, grain size fractions (gravel, sand, and silt/clay) were determined using wet sieving, and Cu and Fe were measured using an atomic adsorption spectrometer. See manuscript for details.

**S3 Table. Metadata per sample.** Table containing information pertaining to each sample such as when (Sampling Season, Season) and where (Site, Location, Estuary) it was taken. A sample was considered to have a muck characteristic if it exceeded one of the muck thresholds: 10% for total organic matter, 60% for silt/clay fraction, and 75 % for water content [7]. Additionally, a sample was considered to have high copper if it exceeded 65 μg/g [69]. ^a^IRL for Indian River Lagoon, ^b^SLE for St. Lucie Estuary, ^c^TOM stands for total organic matter, ^d^Cu for copper, ^e^LoLo for low TOM/low Cu, ^f^HiHi for high TOM/low Cu, ^g^LoHi for low TOM/low Cu, and ^h^HiLo stands for high TOM/low Cu.

**S4 Table. Average monthly means for canal daily discharges (ft^3^/s).** Data was taken from the *United States Geological Services online database [31] or the **South Florida Water Management District’s DBHYDRO online database [30]. ^a^The regional location each canal/stream was found in, ^b^data from the month before and months during each sampling period and ^c^the entire survey. ^d^The the average streamflow from all fourteen stream/canals during the months before and during each sampling period. IRL stands for Indian River Lagoon.

**S5 Table. Environmental parameters statistical analysis.** ^a^Bold text is associated with testing the overall differences within a category with Kruskal-Wallis. ^b^Regular text is associated with pair-wise Dunn testing. ^c^BH stands for Benjamini-Hochberg, ^d^IRL stands for Indian River Lagoon and ^e^SLE stands for St. Lucie Estuary.

**S6 Table. Shannon diversity statistical analysis.** ^a^Bold text is associated with testing the overall differences within a category with Kruskal-Wallis or Mann-Whitney use with a * indicating the latter was used. ^b^Regular text is associated with pair-wise Dunn testing. ^c^BH stands for Benjami-Hochberg, ^d^TOM for total organic matter, ^e^Cu for copper, ^f^IRL for Indian River Lagoon and ^g^SLE for St. Lucie Estuary.

**S7 Table. Full permutational analysis of variance results.** ^a^Bold text is associated with testing the overall differences within a category and ^b^regular text is associated with pair-wise testing. ^c^P(perms) stands for permutational p value, P(MC) for Monte-Carlo p value, ^e^TOM for total organic matter, ^f^Cu for copper, ^g^IRL stands for Indian River Lagoon and ^h^SLE for St. Lucie Estuary.

**S8 Table. Distance-based linear model results.** ^a^Marginal statistical tests are displayed for all variables and ^b^sequential statistical results are shown only if the variable was determined to contribute a statistically significant amount of variation between microbial samples (p value < 0.05). ^c^SS for sum of squares and ^d^AICc stands for An Information Criterion.

## References

1. Indian River Lagoon National Estuary Program [IRLNEP]. Indian River Lagoon Comprehensive Conservation and Mangaement Plan. 2019.

2. East Central Florida Regional Planning Council [ECFRPC], Treasure Coast Regional Planning Council [TCRPC]. Indian River Lagoon Economic Valuation Update [Internet]. 2016. Available from: http://tcrpc.org/special_projects/IRL_Econ_Valu/FinalReportIRL08_26_2016.pdf

3. McKeon CS, Tunberg BG, Johnston CA, Barshis DJ. Ecological drivers and habitat associations of estuarine bivalves. PeerJ [Internet]. 2015;3:e1348. Available from: https://peerj.com/articles/1348

4. Indian River Lagoon National Estuary Program [IRLNEP], St. Johns Water Management District [SJRWMD]. Indian River Lagoon: An Introduction to a National Treasure. 2007.

5. Sun MY, Dafforn KA, Brown M V., Johnston EL. Bacterial communities are sensitive indicators of contaminant stress. Mar Pollut Bull. 2012;64(5):1029–38.

6. Yang Y, He Z, Wang Y, Fan J, Liang Z, Stoffella PJ. Dissolved organic matter in relation to nutrients (N and P) and heavy metals in surface runoff water as affected by temporal variation and land uses–A case study from Indian River Area, south Florida, USA. Agric water Manag. 2013;118:38–49.

7. Trefry JH, Windsor JG, Trocine RP. Toxic Substances in the Indian River Lagoon: Results from the 2006/07 (TOX 2) Survey Contract SJ47613. 2008.

8. St. Johns River Water Management District [SJRWMD]. The Eau Gallie River and Elbow Creek Restoration Dredging Project [Internet]. 2019 [cited 2017 May 3]. Available from: http://www.sjrwmd.com/EGRET/

9. Zhang MK, He ZL, Stoffella PJ, Calvert DV, Yang X, Sime PL. Concentrations and solubility of heavy metals in muck sediments from the St. Lucie Estuary, U.S.A. Environ Geol. 2003;44:1–7.

10. Crump BC, Adams HE, Hobbie JE, Kling GW. Biogeography of bacterioplankton in lakes and streams of an arctic tundra catchment. Ecology. 2007;88(6):1365–78.

11. Diaz RJ, Rosenberg R. Spreading dead zones and consequences for marine ecosystems. Science (80-) [Internet]. 2008;321(5891):926–9. Available from: http://www.ncbi.nlm.nih.gov/pubmed/18703733

12. Johnson JM, Wawrik B, Isom C, Boling WB, Callaghan A V. Interrogation of Chesapeake Bay sediment microbial communities for intrinsic alkane-utilizing potential under anaerobic conditions. FEMS Microbiol Ecol. 2015;

13. Nogales B, Lanfranconi MP, Piña-Villalonga JM, Bosch R. Anthropogenic perturbations in marine microbial communities. Vol. 35, FEMS Microbiology Reviews. 2011. p. 275–98.

14. He ZL, Zhang M, Stoffella PJ, Yang XE. Vertical distribution and water solubility of phosphorus and heavy metals in sediments of the St. Lucie Estuary, South Florida, USA. Environ Geol. 2006;

15. Trefry JH, Trocine RP. Metals in sediments and clams from the Indian River Lagoon, Florida: 2006-7 versus 1992. Florida Sci. 2011;74:43–62.

16. Soule DF, Oguri M, Jones BH. The marine environment of Marina Del Rey: October 1989 to September 1990. Marine Studies of San Pedro Bay, California, Part 20F. Los Angeles; 1991.

17. McMahon PJT. The impact of marinas on water quality. Water Sci Technol. 1989;21(2):39–43.

18. Gadd GM. Metals, minerals and microbes: Geomicrobiology and bioremediation. Vol. 156, Microbiology. 2010. p. 609–43.

19. Magalhães CM, Machado A, Matos P, Bordalo AA. Impact of copper on the diversity, abundance and transcription of nitrite and nitrous oxide reductase genes in an urban European estuary. FEMS Microbiol Ecol. 2011;

20. Cornall A, Rose A, Streten C, Mcguinness K, Parry D, Gibb K. Molecular screening of microbial communities for candidate indicators of multiple metal impacts in marine sediments from northern Australia. Environ Toxicol Chem. 2016;35(2):468–84.

21. Sun MY, Dafforn KA, Johnston EL, Brown M V. Core sediment bacteria drive community response to anthropogenic contamination over multiple environmental gradients. Environ Microbiol. 2013;15(9):2517–31.

22. Vishnivetskaya TA, Mosher JJ, Palumbo A V., Yang ZK, Podar M, Brown SD, et al. Mercury and other heavy metals influence bacterial community structure in contaminated Tennessee streams. Appl Environ Microbiol. 2011;77(1):302–11.

23. Besaury L, Ghiglione JF, Quillet L. Abundance, Activity, and Diversity of Archaeal and Bacterial Communities in Both Uncontaminated and Highly Copper-Contaminated Marine Sediments. Mar Biotechnol. 2014;

24. Garland JL. Potential Extent of Bacterial Biodiversity in the Indian-River Lagoon. Bull Mar Sci. 1995;57(1):79–83.

25. Oni OE, Schmidt F, Miyatake T, Kasten S, Witt M, Hinrichs KU, et al. Microbial communities and organic matter composition in surface and subsurface sediments of the Helgoland mud area, North Sea. Front Microbiol. 2015;

26. Hunter EM, Mills HJ, Kostka JE. Microbial community diversity associated with carbon and nitrogen cycling in permeable shelf sediments. Appl Environ Microbiol. 2006;

27. St. Johns Water Management District [SJRWMD]. Continuous Sensor-based Water Quality Data [Internet]. 2020 [cited 2020 Apr 2]. Available from: http://webapub.sjrwmd.com/agws10/hdswq/

28. Harbor Branch Oceanographic Institute [HBOI]. FAU Harbor Branch Indian River Lagoon Observator [Internet]. 2020 [cited 2020 Apr 2]. Available from: http://fau.loboviz.com/

29. National Weather Service [NWS]. NOWData - NOAA Online Weather Data (Melbourne, FL) [Internet]. 2020 [cited 2020 Apr 2]. Available from: https://w2.weather.gov/climate/xmacis.php?wfo=mlb

30. South Florida Water Management District [SFWMD]. DBHYDRO [Internet]. 2020 [cited 2020 Apr 2]. Available from: https://my.sfwmd.gov/dbhydroplsql/show_dbkey_info.main_menu

31. United States Gelogical Service [USGS]. Current Conditions for Florida: Streamflow [Internet]. 2020 [cited 2020 Apr 2]. Available from: https://waterdata.usgs.gov/fl/nwis/current/?type=flow&group_key=basin_cd

32. Munksgaard NC, Parry DL. Trace metals, arsenic and lead isotopes in dissolved and particulate phases of North Australian coastal and estuarine seawater. Mar Chem. 2001;75(3):165–84.

33. Cornall AM, Beyer S, Rose A, Streten-Joyce C, McGuinness K, Parry D, et al. HCl-Extractable Metal Profiles Correlate with Bacterial Population Shifts in Metal-Impacted Anoxic Coastal Sediment from the Wet/Dry Tropics. Geomicrobiol J [Internet]. 2013;30(1):48–60. Available from: http://www.tandfonline.com/doi/abs/10.1080/01490451.2011.653083

34. Caporaso JG, Lauber CL, Walters W a, Berg-Lyons D, Huntley J, Fierer N, et al. Ultra-high-throughput microbial community analysis on the Illumina HiSeq and MiSeq platforms. ISME J [Internet]. 2012;6(8):1621–4. Available from: http://dx.doi.org/10.1038/ismej.2012.8

35. Earth Microbiome Project [EMP]. “16S rRNA Amplification Protocol” [Internet]. 2019 [cited 2017 May 3]. Available from: http://www.earthmicrobiome.org/emp-standard-protocols/16s/

36. Andrews S. FastQC. Babraham Bioinforma. 2010;

37. Krueger F. Trim Galore [Internet]. Babraham Bioinformatics. 2016. Available from: https://github.com/FelixKrueger/TrimGalore

38. Bolyen E, Rideout JR, Dillon MR, Bokulich NA, Abnet CC, Al-Ghalith GA, et al. Reproducible, interactive, scalable and extensible microbiome data science using QIIME 2. Nature Biotechnology. 2019.

39. QIIME 2 Development Team. “Moving Pictures” tutorial [Internet]. 2019 [cited 2019 Mar 6]. Available from: https://docs.qiime2.org/2018.11/tutorials/moving-pictures/

40. Köster J, Rahmann S. Snakemake-a scalable bioinformatics workflow engine. Bioinformatics. 2012;

41. Rognes T, Flouri T, Nichols B, Quince C, Mahé F. VSEARCH: a versatile open source tool for metagenomics. PeerJ. 2016;

42. Amir A, McDonald D, Navas-Molina JA, Kopylova E, Morton JT, Zech Xu Z, et al. Deblur Rapidly Resolves Single-Nucleotide Community Sequence Patterns. mSystems. 2017;

43. Klindworth A, Pruesse E, Schweer T, Peplies J, Quast C, Horn M, et al. Evaluation of general 16S ribosomal RNA gene PCR primers for classical and next-generation sequencing-based diversity studies. Nucleic Acids Res. 2013;41(1).

44. Pedregosa F, Varoquaux G, Gramfort A, Michel V, Thirion B, Grisel O, et al. Scikit-learn: Machine learning in Python. J Mach Learn Res. 2011;

45. Katoh K, Standley DM. MAFFT multiple sequence alignment software version 7: improvements in performance and usability. Mol Biol Evol. 2013;

46. Lane DJ. Nucleic Acid Techniques in Bacterial Systematics. In: Stackebrandt E, Goodfellow M, editors. New York: John Wiley and Sons; 1991. p. 115–75.

47. Price MN, Dehal PS, Arkin AP. FastTree 2 - Approximately maximum-likelihood trees for large alignments. PLoS One. 2010;

48. Bradshaw DJ. BioProject: PRJNA594146 [Internet]. 2020. Available from: https://dataview.ncbi.nlm.nih.gov/object/PRJNA594146?reviewer=j30a930a8h6kh633mp24p9c42p

49. RStudio Team. RStudio: Integrated Development for R. [Online] RStudio, Inc., Boston, MA URL http://www.rstudio.com. 2015.

50. R Development Core Team. Computational Many-Particle Physics. R Found Stat Comput [Internet]. 2008;739. Available from: http://link.springer.com/10.1007/978-3-540-74686-7

51. McMurdie PJ, Holmes S. Phyloseq: An R Package for Reproducible Interactive Analysis and Graphics of Microbiome Census Data. PLoS One. 2013;8(4).

52. Oksanen J, Blanchet FG, Kindt R, Legendre P, McGlin D, Minchin PR, et al. vegan: Community Ecology Package [Internet]. 2019. Available from: https://cran.r-project.org/package=vegan

53. Wickham H. ggplot2: elegant graphics for data analysis. Journal of the Royal Statistical Society: Series A (Statistics in Society). 2016.

54. Wickham H. Reshaping data with the reshape package. J Stat Softw. 2007;

55. Wickham H. tidyverse: Easily Install and Load the “Tidyverse.” R package version 1.2.1. 2017.

56. Ogle D, Wheeler P, Dinno A. FSA: Fisheries Stock Analysis [Internet]. 2019. Available from: https://github.com/droglenc/FSA

57. Bolyen E, Rideout JR, Dillon MR, Bokulich NA, Abnet C, Ghalith GA Al, et al. QIIME 2: Reproducible, interactive, scalable, and extensible microbiome data science. PeerJ Prepr. 2018;

58. Ziegler M, Seneca FO, Yum LK, Palumbi SR, Voolstra CR. Bacterial community dynamics are linked to patterns of coral heat tolerance. Nat Commun. 2017;

59. Anderson MJ, Gorley RN, Clarke KR. PERMANOVA+ for PRIMER: Guide to Software and Statistical Methods. Plymouth, UK. 2008.

60. Gutiérrez NL, Hilborn R, Defeo O, Clarke KR, Gorley RN, Barrientos IM, et al. Getting started with PRIMER v7 Plymouth Routines In Multivariate Ecological Research. Rev Mex Biodivers. 2018;

61. Clarke KR, Gorley RN. ’PRIMER v7: User Manual / Tutorial. 2015.

62. Ward L, Taylor MW, Power JF, Scott BJ, McDonald IR, Stott MB. Microbial community dynamics in Inferno Crater Lake, a thermally fluctuating geothermal spring. ISME J. 2017;

63. Behera P, Mahapatra S, Mohapatra M, Kim JY, Adhya TK, Raina V, et al. Salinity and macrophyte drive the biogeography of the sedimentary bacterial communities in a brackish water tropical coastal lagoon. Sci Total Environ. 2017;

64. Gooch JW. Kruskal-Wallis Test. In: Encyclopedic Dictionary of Polymers. 2011.

65. Gooch JW. Mann-Whitney U Test. In: Encyclopedic Dictionary of Polymers. 2011.

66. Elliott AC, Hynan LS. A SAS^®^ macro implementation of a multiple comparison post hoc test for a Kruskal-Wallis analysis. Comput Methods Programs Biomed. 2011;

67. Benjamini Y, Hochberg Y. Controlling the False Discovery Rate: A Practical and Powerful Approach to Multiple Testing. J R Stat Soc Ser B. 1995;

68. Bradshaw DJ. General_16S_Amplicon_Sequencing_Analysis [Internet]. 2019 [cited 2019 Feb 12]. Available from: https://github.com/djbradshaw2/General_16S_Amplicon_Sequencing_Analysis

69. Australian and New Zealand Environment and Conservation Council [ANZECC]. Australian and New Zealand guidelines for fresh and marine water quality. A. 2000.

70. Sime P. St. Lucie Estuary and Indian River Lagoon conceptual ecological model. Wetlands. 2006;

71. Trefry JH, Chen N, Trocine RP, Metz S. Impingement of Organic-Rich, Contaminated Sediments on Manatee Pocket, Florida. Florida Sci. 1992;160–71.

72. Badylak S, Phlips EJ. Spatial and temporal patterns of phytoplankton composition in a subtropical coastal lagoon, the Indian River Lagoon, Florida, USA. J Plankton Res. 2004;

73. United States Environmental Protection Agency [USEPA]. National Management Measures Guidance to Control Nonpoint Source Pollution from Marinas and Recreational Boating, EPA 841-B-01-005. 2001.

74. Dell’Anno A, Beolchini F, Rocchetti L, Luna GM, Danovaro R. High bacterial biodiversity increases degradation performance of hydrocarbons during bioremediation of contaminated harbor marine sediments. Environ Pollut. 2012;

75. Mohit V, Archambault P, Lovejoy C. Resilience and adjustments of surface sediment bacterial communities in an enclosed shallow coastal lagoon, Magdalen Islands, Gulf of St. Lawrence, Canada. FEMS Microbiol Ecol. 2015;

76. Quero GM, Perini L, Pesole G, Manzari C, Lionetti C, Bastianini M, et al. Seasonal rather than spatial variability drives planktonic and benthic bacterial diversity in a microtidal lagoon and the adjacent open sea. Mol Ecol. 2017;

77. Guttman DS, McHardy AC, Schulze-Lefert P. Microbial genome-enabled insights into plant-microorganism interactions. Nature Reviews Genetics. 2014.

78. Brenner DJ, Krieg NR, Staley JT, editors. Bergey’s Manual of Systematic Bacteriology - Vol 2: The Proteobacteria Part C -The Alpha-, Beta-, Delta-, and Epsilonproteobacteria. 2nd ed. New York: Springer Science+Business Media, Inc.; 2005.

79. Du ZJ, Wang ZJ, Zhao JX, Chen GJ. Woeseia oceani gen. Nov., sp. nov., a chemoheterotrophic member of the order Chromatiales, and proposal of Woeseiaceae fam. nov. Int J Syst Evol Microbiol. 2016;

80. Mußmann M, Pjevac P, Krüger K, Dyksma S. Genomic repertoire of the Woeseiaceae/JTB255, cosmopolitan and abundant core members of microbial communities in marine sediments. ISME J. 2017;

81. Spring S, Scheuner C, Göker M, Klenk HP. A taxonomic framework for emerging groups of ecologically important marine gammaproteobacteria based on the reconstruction of evolutionary relationships using genome-scale data. Front Microbiol. 2015;

82. Park S, Yoshizawa S, Inomata K, Kogure K, Yokota A. Halioglobus japonicus gen. nov., sp. nov. and halioglobus pacificus sp. nov., members of the class gammaproteobacteria isolated from seawater. Int J Syst Evol Microbiol. 2012;

83. Krieg NR, Staley JT, Brown DR, Hedlund BP, Paster BJ, Ward NL, et al., editors. Bergey’s Volume 4 Bacteroidetes, Acidobacteria, Spirochaetes, Tenericutes (Mollicutes), Acidobacteria, Fibrobacteres, Fusobacteria, Dictyoglomi, Gemmatimonadetes, Lentisphaerae, Verrucomicrobia, Chlamydiae, and Planctomycetes. 2nd ed. New; 2011.

84. McIlroy SJ, Kirkegaard RH, Dueholm MS, Fernando E, Karst SM, Albertsen M, et al. Culture-independent analyses reveal novel anaerolineaceae as abundant primary fermenters in anaerobic digesters treating waste activated sludge. Front Microbiol. 2017;

85. Campbell BJ, Yu L, Heidelberg JF, Kirchman DL. Activity of abundant and rare bacteria in a coastal ocean. Proc Natl Acad Sci. 2011;

86. Gaidos E, Rusch A, Ilardo M. Ribosomal tag pyrosequencing of DNA and RNA from benthic coral reef microbiota: Community spatial structure, rare members and nitrogen-cycling guilds. Environ Microbiol. 2011;

87. Gillan DC, Danis B, Pernet P, Joly G, Dubois P. Structure of sediment-associated microbial communities along a heavy-metal contamination gradient in the marine environment. Appl Environ Microbiol. 2005;71(2):679–90.

88. Oppenheimer CH. Bacterial activity in sediments of shallow marine bays. Geochim Cosmochim Acta. 2003;

89. Mahmoudi N, Robeson MS, Castro HF, Fortney JL, Techtmann SM, Joyner DC, et al. Microbial community composition and diversity in Caspian Sea sediments. FEMS Microbiol Ecol. 2015;91(1).

90. Waite DW, Vanwonterghem I, Rinke C, Parks DH, Zhang Y, Takai K, et al. Comparative genomic analysis of the class Epsilonproteobacteria and proposed reclassification to epsilonbacteraeota (phyl. nov.). Front Microbiol. 2017;

91. Hou D, Zhang P, Zhang J, Zhou Y, Yang Y, Mao Q, et al. Spatial variation of sediment bacterial community in an acid mine drainage contaminated area and surrounding river basin. J Environ Manage. 2019;

92. Fernández-Cadena JC, Ruíz-Fernández PS, Fernández-Ronquillo TE, Díez B, Trefault N, Andrade S, et al. Detection of sentinel bacteria in mangrove sediments contaminated with heavy metals. Mar Pollut Bull. 2020;

93. Inagaki F, Takai K, Nealson KH, Horikoshi K. Sulfurovum lithotrophicum gen. nov., sp. nov., a novel sulfur-oxidizing chemolithoautotroph within the E-Proteobacteria isolated from Okinawa Trough hydrothermal sediments. Int J Syst Evol Microbiol. 2004;

94. Mohr KI, Garcia RO, Gerth K, Irschik H, Müller R. Sandaracinus amylolyticus gen. nov., sp. nov., a starch-degrading soil myxobacterium, and description of Sandaracinaceae fam. nov. Int J Syst Evol Microbiol. 2012;

95. Schlesner H, Rensmann C, Tindall BJ, Gade D, Rabus R, Pfeiffer S, et al. Taxonomic heterogeneity within the Planctomycetales as derived by DNA-DNA hybridization, description of Rhodopirellula baltica gen. nov., sp. nov., transfer of Perillula marina to the genus Blastopirellula gen. nov. as Blastopirellula marina comb. nov. an. Int J Syst Evol Microbiol. 2004;

96. Bondoso J, Albuquerque L, Nobre MF, Lobo-da-Cunha A, da Costa MS, Lage OM. Roseimaritima ulvae gen. nov., sp. nov. and Rubripirellula obstinata gen. nov., sp. nov. two novel planctomycetes isolated from the epiphytic community of macroalgae. Syst Appl Microbiol. 2015;

